# GDF15 reprograms the microenvironment to drive the development of uveal melanoma liver metastases

**DOI:** 10.1101/2025.05.07.652654

**Authors:** Sathya Neelature Sriramareddy, Navya Siddarajappa, Jun Sung Park, Thanh Nguyen, Jeffim N. Kuznetsoff, Stefan Kurtenbach, James J. Dollar, David J. Adams, Jonathan D. Licht, Zelia M. Correa, Y. Ann Chen, J. William Harbour, Keiran S.M. Smalley

## Abstract

Uveal melanoma (UM) results in fatal liver metastasis, yet little is known about the interactions between UM and host cells in the tumor microenvironment that promote this distinctive proclivity. Here, we used single cell (sc)-RNA-Seq analysis of UM-hepatic stellate cell (HSC) co-cultures to demonstrate that HSCs enriched for UM cell states that expressed genes implicated in cell survival, metabolic reprogramming and angiogenesis. A lead candidate driver of HSC reprogramming was the TGF-β family member GDF15, which was associated with a metastatic UM phenotype. Silencing of BAP1 in UM cells led to increased GDF15 expression and accumulation of H3K27ac marks at the GDF15 promoter. Treatment of HSCs with GDF15 led to increased expression extracellular matrix proteins, inflammatory cytokines and angiogenic factors, including IL-8. Both exogenous GDF15, IL-8 and conditioned media from UM-HSC co-cultures increased endothelial cell network formation *in vitro,* an effect that was blocked by anti-GDF15 antibodies. In multiple models of metastatic UM, silencing of GDF15 inhibited the outgrowth of metastatic lesions, associated with reduced deposition of extracellular matrix and recruitment of endothelial cells. UM liver metastasis development is dependent upon GDF15-mediated remodeling of the liver microenvironment leading to an angiogenic response and matrix deposition that supports tumor growth.

## Introduction

Uveal melanoma (UM) is a highly aggressive tumor derived from melanocytes residing in the uveal tract of the eye. Although disseminated disease is not frequently observed at the time of diagnosis, approximately half of patients eventually succumb to metastases, often many years after successful treatment of the primary tumor^1^. The major site for UM metastasis is the liver. Patients are stratified into groups with low (Class 1) vs. high (Class 2) risk of metastasis on the basis of a 15 gene expression profile (GEP) assay^2^ performed on the primary tumor prior to treatment. Class 1 tumors have a more melanocyte-like differentiation pattern and are associated with a low risk of metastatic dissemination^3^. By contrast, Class 2 tumors have a less differentiated transcriptional profile and a high risk of metastatic dissemination of 70-80% at 5 years^3^. This classification was recently improved by the inclusion of the expression of the cancer-testis antigen Preferentially Expressed Antigen in Melanoma (PRAME), which was associated with increased metastatic risk in both Class 1 and 2 tumors ^4, 5^. The most frequent driver oncogene in UM are mutations in the small G-proteins GNAQ or GNA11 which activate the mitogen activated protein kinase (MAPK) pathway ^6, 7^. GNAQ/GNA11 mutations can be found in benign nevi, suggesting that additional genetics events are required for tumor progression ^6–8^. One of the major genetic drivers of the Class 2 phenotype is loss or inactivating mutations in the H2A ubiquitin hydrolase BAP1^9^. BAP1 is the catalytic subunit of the Polycomb repressive deubiquitinase (PR-DUB) complex that deubiquitinates histone H2A modulating gene expression ^10^. Bi-allelic inactivation of BAP1 is associated with global epigenetic reprogramming resulting in a dedifferentiated, migratory phenotype associated with a high rate (>70%) of liver metastasis^2^. PRAME is a CUL2 ubiquitin ligase subunit whose aberrant expression in UM has been associated with genomic instability resulting from DNA damage, aneuploidy and telomere dysfunction^11^.

Little is known about the cellular microenvironment of UM liver metastases. In other liver tumors such as hepatocellular carcinoma, interactions between cancer cells and hepatic stellate cells (HSCs) are a key driver of tumor progression^12^. In healthy liver HSCs comprise only 5-8% of the total cells. Upon injury or inflammation, HSC numbers increase, and they adopt an activated, myofibroblast-like phenotype^13, 14^. Prior *in vitro* studies have shown that HSCs may contribute to escape of UM cells from targeted therapies including MEK inhibitors and bromodomain inhibitors^15, 16^. HSCs are also a potential source of growth factors and cytokines that stimulate UM cells^17, 18^. Nevertheless, the role of HSCs in the initiation and progression of UM liver metastases remains poorly characterized.

In the present study we explored the mechanisms by which UM cells metastasize to the liver. We found that the UM cell interaction with HSCs shapes the transcriptional profile of the UM cells, increasing the expression of genes involved in cell survival, inflammatory programs and a growth differentiation factor (GDF)-15-mediated angiogenic program. GDF15 activates HSCs increasing expression of pro-angiogenic IL-8, which along with GDF15 itself, drives formation of vascular networks. Silencing of GDF15 in the UM cells abrogated the formation of liver metastases. Together these studies demonstrate a key role for UM-derived GDF15 in mediating crosstalk with HSCs and endothelial cells, facilitating the development of liver metastases.

## Results

### HSCs alter the transcriptional state of UM cells

To gain insights into the cell-cell communication between HSCs and UM cells we grew liver metastasizing MP41 UM cells in either monoculture or in co-culture with the LX2 HSC cells for 48 hours before undertaking scRNA-Seq analysis. The two cell populations were easily distinguished by their expression of melanocytic lineage markers (*PMEL*, *MITF*) and those known to be expressed in HSCs (e.g. *COL1A1*, *VCL*, *SPARC*, *PDGFA*) (**Figure 1A**). These analyses revealed multiple transcriptional states within the UM cell population (**Figure 1B, Supplemental Figure 1**). In the presence of the HSCs, there was a marked change in the transcriptional profile of the UM cells, and particularly an enrichment for transcriptional state #8, and a simultaneous decrease in the prevalence of state #1 (**Figure 1B**). We next used STRING analysis to build networks of genes enriched in UM cells grown in monoculture and co-culture. It was found that upon co-culture the presence of the HSCs enriched for genes in the UM cells associated with increase survival, altered metabolic states and increased angiogenic responses (**Figure 1C**). Multiple genes were identified in transcriptional state #8 including many involved in UM-specific cell signaling (*RASGRP3, MITF*), cell survival (*BIRC7*), transcriptional regulation and cell-cell communication (*GDF15*) (**Figures 1C,D**) ^19, 20^.

**Figure 1:**
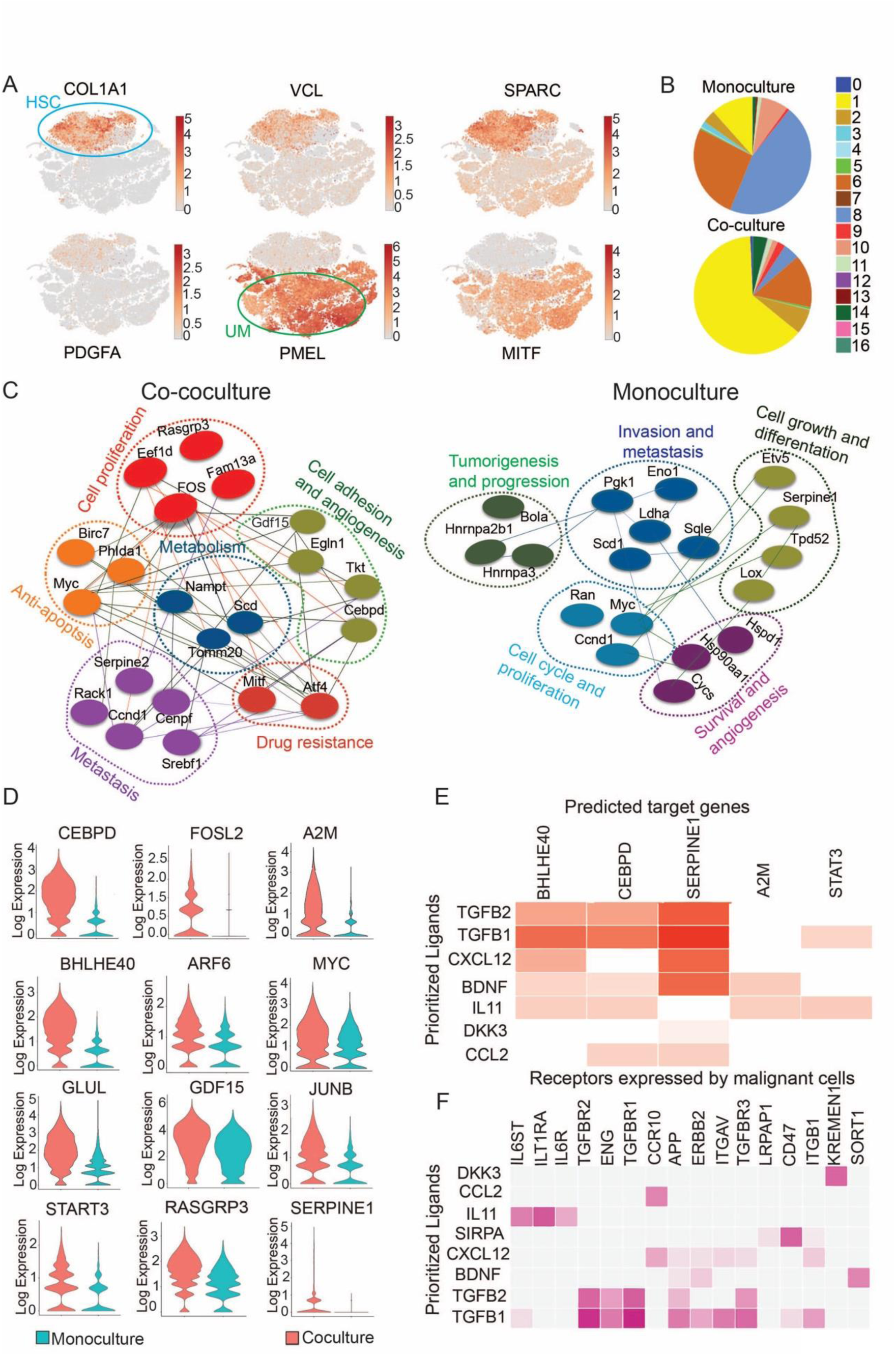
Hepatic stellate cells (HSCs) modulate the transcriptional state of UM cells. **A:** t-SNE plots from UM (MP41) and HSCs (LX2) grown in monoculture and co-culture, indicating key lineage genes in HSCs (*COL1A1, VCL, SPARC, PDGFA*) and UM cells (*PMEL, MITF*). Scale bar shows log normalized gene expression. **B:** Unbiased clustering of cell states from UM cells grown as a monoculture or co-culture identifies multiple transcriptional states that change upon co-culture with HSCs. **C:** STRING analysis of differentially expressed genes in the UM cells between monoculture and co-culture. **D:** Violin plot of differentially expressed genes in the UM cells between the monoculture and co-culture. **E:** NicheNet analysis of activated receptors->differentially expressed genes in D. **F:** NicheNet analysis of inferred receptor-ligand pairs based upon analysis in E.

NicheNet was utilized to identify potential regulators of cell-cell communication between the HSCs and UM cells. We first determined the most significant differentially expressed genes in the UM cells following co-culture, and then identified the ligand-receptor pairs likely to be responsible for these changes (**Figures 1E,F**). *TGFB1, TGFB2, CXCL12, BDNF, IL11, DKK3* and *CCL2* were expressed in the HSCs, and the growth factors and cytokines encoded by these genes were predicted to alter expression of genes differentially expressed upon co-culture of UM cells with HSC, including *STAT3, A2M, BHLHE40, SERPINE1* and *CEBPD* (**Figure 1D**), through multiple receptors expressed on the UM cells (**Figure 1F**). Together these data suggest that communication from HSCs to UM cells alters patterns of gene expression in UM cells, stimulating pathways known to increase tumor cell survival.

### Metastatic UM cells rewire their cell-cell contacts to communicate with hepatic stellate cells and endothelial cells

We next examined the microenvironment of primary UM and UM liver metastases through CellChat analysis of our previously published single cell RNA-seq UM dataset ^21^. This analysis indicated that in primary UM the two major drivers of cell-cell communication were the fibroblasts and the endothelial cells, which mostly impacted each other, as well as immune cells (CD8+, CD4+, NK cells and monocytes) (**Figure 2A**). In the UM liver metastases, the HSCs were a predominant driver of cell-cell communication, with interactions between HSCs and endothelial cells, CD8+ T cells, CD4+ T cells and NK cells being observed (**Figure 2B**). The UM cells exhibited increased interactions with CD8+ T cells and the HSCs. The patterns of incoming and outgoing signaling in the UM metastases revealed a marked increase in outgoing signals from the HSCs and endothelial cells compared to the extent of signaling in the primary UM (**Figures 2C,D**). An analysis of the gene expression in the tumor microenvironment of primary UM identified four signaling patterns. The first, shared by endothelial cells and B cells, was associated with adhesion signaling (*PECAM, Laminin, EPHA* etc) (**Figure 2E**). Pattern 2 was specific to lymphocytes (CD4+, CD8+ and NK cells) and centered on immune recognition and adhesion (*NOTCH, CLEC, CD45* and *CD99*). Pattern three was associated with monocytes and involved a variety of adhesion molecules and growth factors involved in inflammation (*TNF, ICAM, GAS, CCL)*. Pattern 4 was noted in tumor cells and fibroblasts and involved signals involved in ECM and adhesion (*COLAI, FN1, EPHB*) as well as growth factors (*EDN, PTN, MK*) (**Figure 2E**).

**Figure 2:**
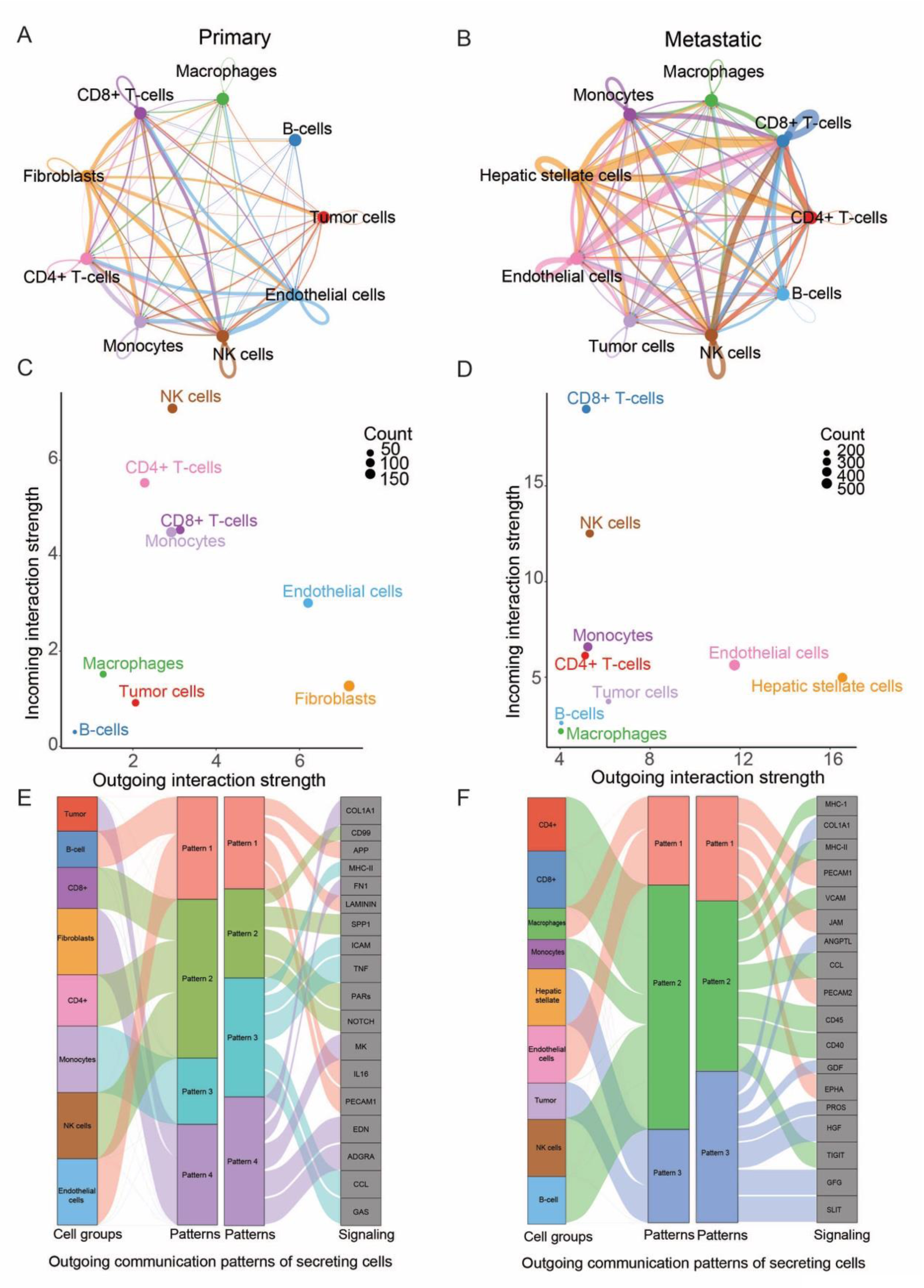
Human metastatic uveal melanoma cells communicate with endothelial cells and hepatic stellate cells. **A**: CellChat analysis showing the number and strength of cell-cell interactions in primary UM. **B**: CellChat analysis showing the number and strength of cell-cell interactions in metastatic UM. **C:** The total numbers of incoming and outgoing cell-cell interactions in primary UM samples. **D:** The total numbers of incoming and outgoing cell-cell interactions in primary UM samples. **E**: Identified patterns of cell-cell signaling and the genes associated with each signaling pattern for the primary UM samples through CellChat analysis. **F:** Identified patterns of cell-cell signaling and the genes associated with each signaling pattern for the metastatic UM samples through CellChat analysis.

In the metastases, the UM cells and HSCs shared a common signaling pattern (pattern #3), which was characterized by expression of genes encoding growth factors including *HGF, GDF15, FGF, ANGPTL* and *PROS1*, as well as Collagen (*COL1A1*) (**Figure 2F**). Marked alterations were also noted in the lymphocyte compartment of the metastatic lesions, which all associated with signaling pattern #2, involving T cell exhaustion markers such as *TIGIT*, genes encoding MHC I and II molecules, and markers of endothelial cell adhesion (*VCAM*). Together these data suggested that liver metastatic UM cells had different patterns of cell-cell communication from primary UM cells and that they interacted with host HSCs and endothelial cells.

### Cross-talk between HSCs and UM cells leads to an increase in pro-angiogenic and inflammatory factors

To determine if the interaction of HSCs and UM cells led to the upregulation of growth factors that contributed to the progression of liver metastases we used growth factor arrays to profile the secreted growth factors from two metastatic UM cell lines (MP41 and UM066) grown in monoculture or in co-culture with HSCs (**Figures 3A,B**). The HSCs expressed Endoglin, bFGF, DKK-1, MIF, SDF1 and Emmprin at baseline, whereas the UM cell lines expressed GDF15, ICAM-1, Endoglin, MIF, osteopontin and Emmprin, (**Figures 3A,B**). When the HSCs were co-cultured with either the MP41 or UMM061 UM cells there was an upregulation in multiple growth factors associated with the angiogenic response and inflammation including VEGF, EGF, VCAM-1 and IL-8 (depending on the cell line) (**Figure 3B**). As IL-8 was strongly induced by both UM cell lines, we extended the findings to a larger panel of UM cell lines. Although some (such as 92.1 and MM28) did secrete some IL-8, levels were generally quite low across the majority of UM cell lines and the HSCs grown as monocultures (**Figure 3C**). Upon co-culture, there was a significant increase in IL-8 secretion for all UM cell lines except 92.1 (**Figures 3C,D**). To determine which cell type elicited the key angiogenic factors, we returned to scRNA-Seq data which showed the highest expression of VEGFA, FGF2 and IL-8 in the HSCs and the highest expression of GDF15 and SPP1 (osteopontin) in UM cells (**Figure 3E)**. These data suggested that cell-cell crosstalk between UM and HSCs led to the increased secretion of multiple growth factors involved in inflammation and the angiogenic response.

**Figure 3:**
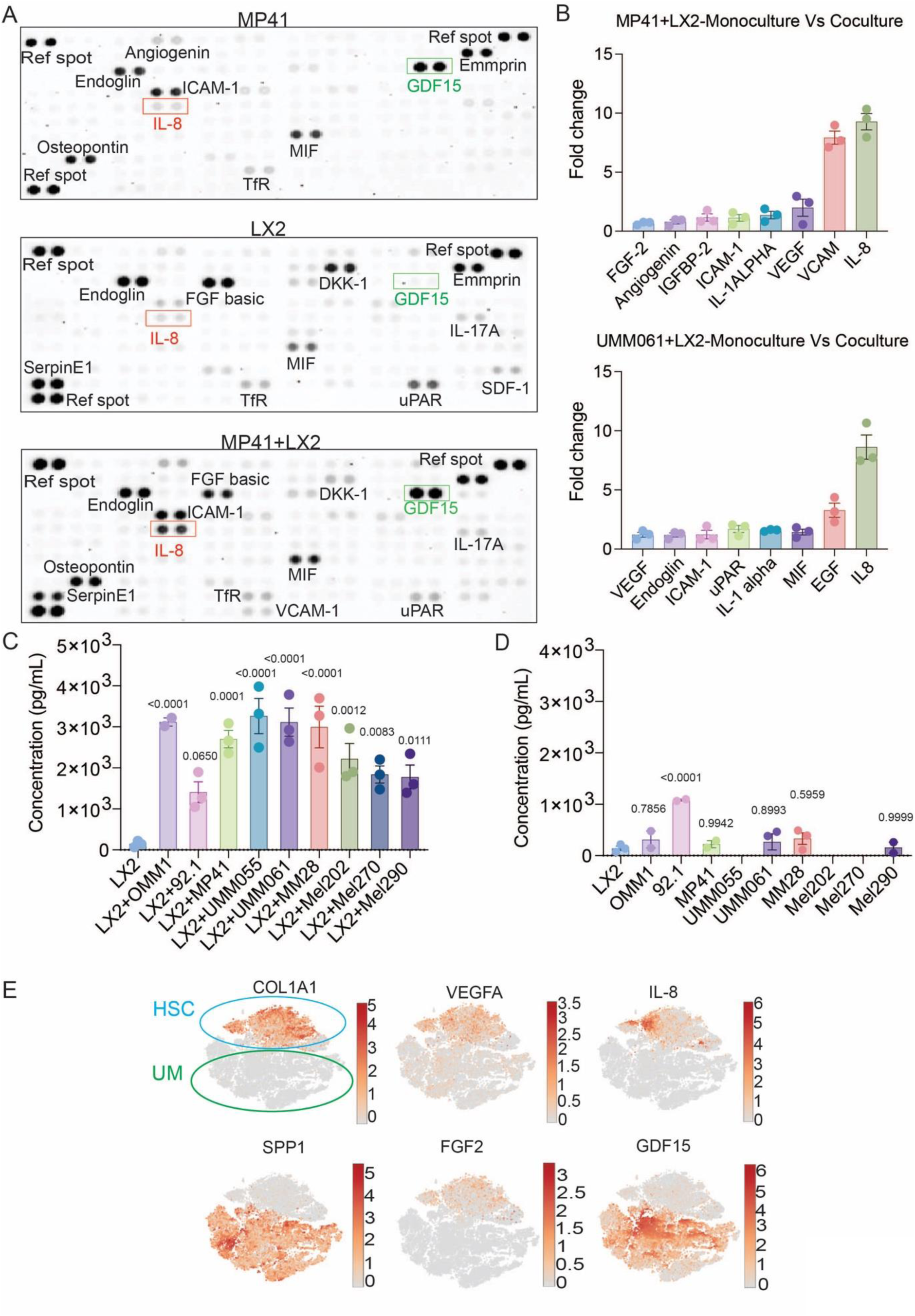
UM-HSC crosstalk is associated with an increased expression of pro-angiogenic factors. **A:** Growth factor arrays of MP41 UM cells and LX2 cells growth in monoculture and MP41-LX2 co-cultures. **B:** Quantification of growth factors increased in MP41 and UM066 UM cells upon co-culture with LX2 cells. **C:** ELISA for IL-8 expression from LX2 cell monocultures and a panel of UM cell lines grown in co-culture with LX2 cells. **D:** ELISA assay showing baseline expression of IL-8 in a panel of UM cell lines. **E:** t-SNE plots of scRNA-seq from UM-HSC co-cultures demonstrating the expression of angiogenic factors in HSCs and not the UM cells. Scale bar shows log normalized gene expression.

### Metastatic UM cells express increased levels of GDF15

As our CellChat analysis of UM cells and the scRNA-seq of UM-HSC co-cultures (**Figures 1-2**) identified GDF15 as a potential driver of UM-HSC communication we next investigated the clinical relevance of this finding. We interrogated scRNA-Seq data from a cohort of 6 individual datasets from primary and metastatic UM and identified UM cells as being the primary source of GDF15 (**Figure 4A**), highlighting the clinical relevance of the cell line mode. Analysis of scRNA-seq data that was classified according to Class 1 and Class 2 UM status ^21^ demonstrated that GDF15 expression was increased in both Class 2 UM samples from both primary and metastatic sites (**Figure 4B**). As both PRAME expression and BAP1 loss are associated with decreased metastasis-free survival ^4^ we next investigated the relationship between PRAME expression, BAP1 status and GDF15 expression. In the cell line panel, increased expression of GDF15 was associated with both expression of PRAME (Mel270, UMM061, MM28 and MP41) and loss of BAP1 (UMM061, MM28) (**Figure 4C**). shRNA-mediated silencing of BAP1 in two UM cell lines, Mel202 and 92.1, was associated with increased GDF15 mRNA expression (**Figure 4D**). ChIP-Seq analysis demonstrated that silencing of BAP1 in Mel202 UM cells led to an increase in H3K27ac at the GDF15 locus, suggesting that BAP1 loss led to an increase in GDF15 transcription (**Figure 4E**). As scRNA-Seq data suggested that UM-HSC co-culture increased GDF15 expression in the UM cells (**Figure 1**), we found that co-culture of a panel of UM cell lines with human primary HSCs in nearly every case led to an increase in GDF15 secretion, even in UM cell lines with high basal GDF15 secretion (**Figure 4F**). These data suggested that UM-HSC crosstalk could potentiate the release of GDF15 from UM cells.

**Figure 4:**
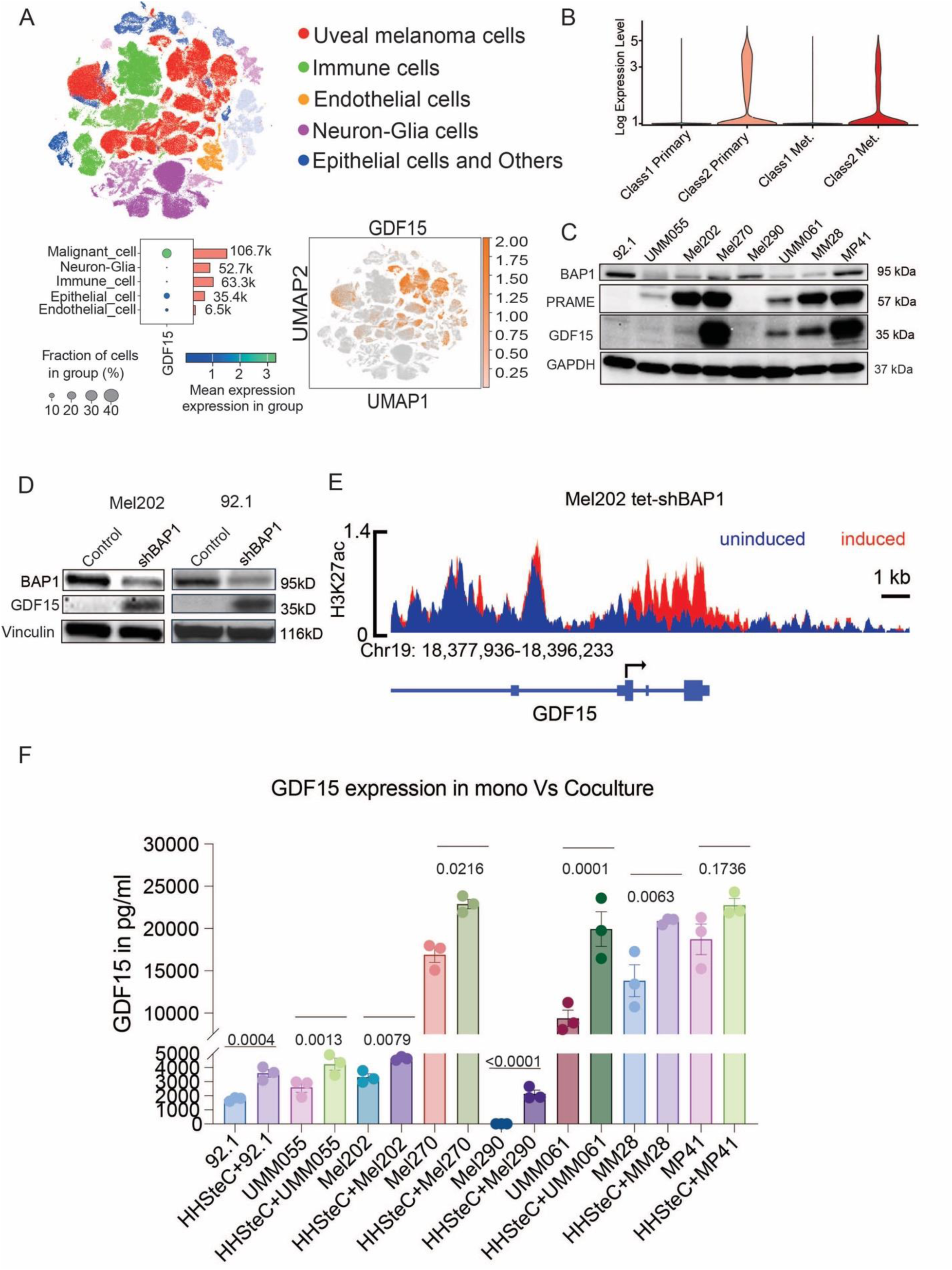
GDF15 expression is upregulated in BAP-1 negative or PRAME expressing UM cells. **A:** UMAP of 6 individual clinical UM scRNA-Seq datasets. Data shows major cell types identified and then GDF15 expression in the UM cells (malignant cells) **B:** Violin plots showing expression of GDF15 in a scRNA-Seq analysis of human Class 1 and Class 2 UM. **C:** Western Blot showing the expression of BAP1, PRAME and GDF15 in a panel of UM cell lines. **D:** Silencing of BAP1 is associated with increased expression of GDF15 in Mel202 and 92.1 UM cells. BAP1 was silenced using siRNA and resulting lysates probed for expression of BAP1 and GDF15. **E:** ChIP-Seq analysis of Mel202 cells expressing BAP1 or silenced for BAP1 demonstrates an increase in H3K27ac at the GDF15 promoter following BAP1 knockdown. **F:** Analysis of GDF15 secretion by ELISA assay demonstrates that co-culture of UM cells with HSCs is associated with increased GDF15 expression, even in UM cell lines with high basal expression of GDF15.

### UM-derived GDF15 leads to increased expression of ECM proteins and angiogenic factors in HSCs

To more comprehensively define the effects of GDF15 upon HSCs, LX2 cells were treated with human recombinant GDF15 for 24 hours followed by RNA-Seq. This showed that GDF15 led to increased expression of multiple genes encoding ECM components (e.g. *COL7A1, COL27A1*) including some implicated in angiogenesis (*VCAM1, ANGPLT4, PDGF-B, FLT1*) (**Figure 5A**). The most enriched pathways included those implicated in hypoxia responses, angiogenesis and immune responses/inflammation (*STAT3, IFNγ, TNFα*) (**Figure 5B**). Treatment of HSCs with GDF15 led to increased phospho-STAT3 expression, an effect accompanied by increased expression of IL-8 (**Figure 5C**). By contrast, a different UM-derived growth factor (osteopontin (SPP1)), also increased STAT3 signaling in HSCs but did not stimulate IL-8 expression (**Figure 5C**). GDF15 and conditioned media from MP41 UM cells increased the expression of known HSC activation markers including *COL1A1* and *PDGFRB* in primary human hepatic stellate cells (HHStec) (**Figure 5D**). Furthermore, conditioned media from MP41 and UMM061 metastatic UM cell lines induced the expression of *COL1A1* in LX2 HSC cells (**Figure 5E**). CellChat analysis of the human UM liver metastasis data confirmed *COL1A1* and *COL1A2* as encoding two ECM proteins that could interact with integrins expressed on UM cells (**Figure 5F**).

**Figure 5:**
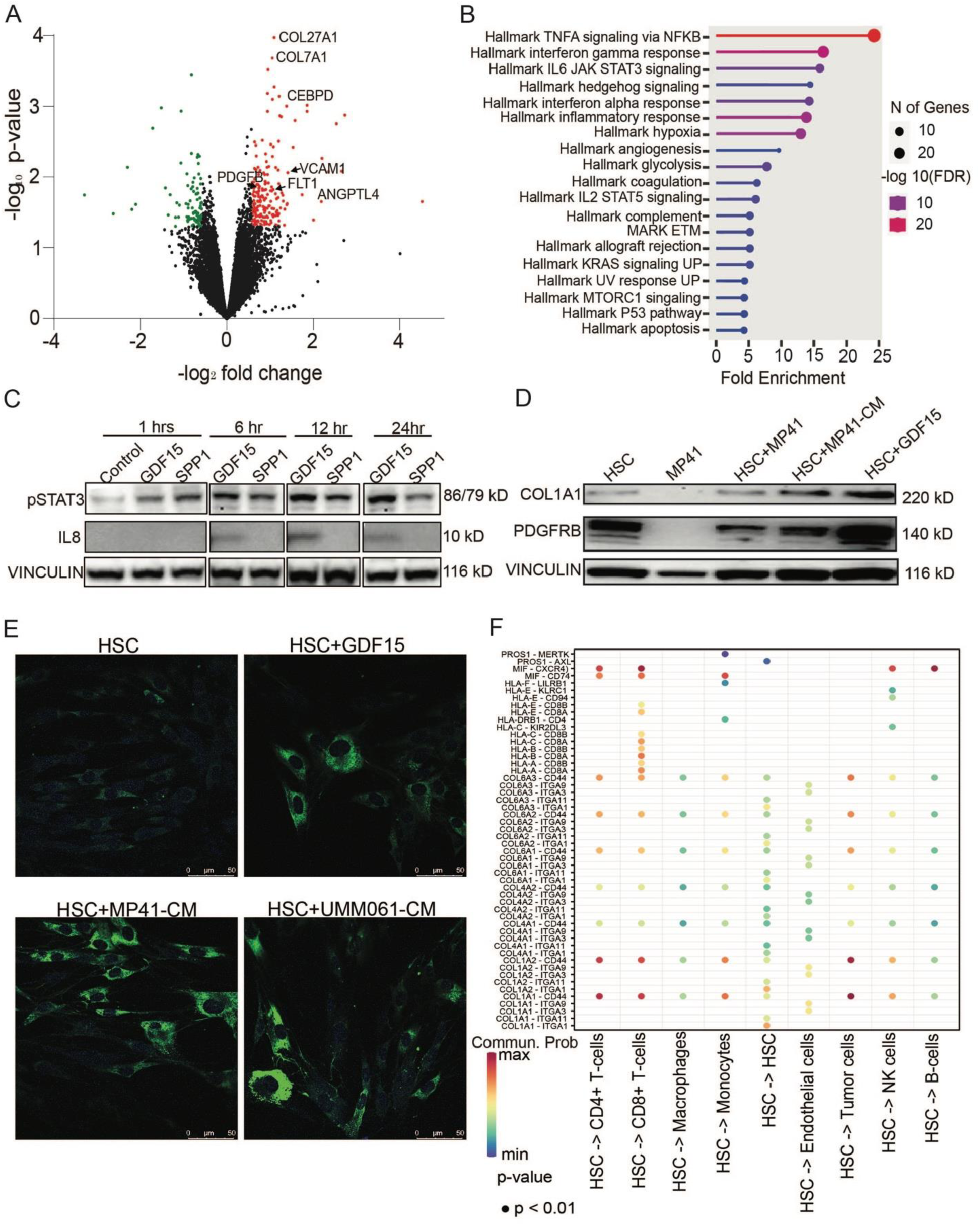
GDF15 drives differentiation and increased growth factor expression in HSCs. **A:** RNA-seq analysis of HSCs (LX2) treated with GDF15 (1μg/ml, 24 hr) identifies increased expression of genes encoding multiple ECM proteins (*COL7A1*), and molecules involved in angiogenesis (*VCAM1, ANGPLT4, PDGFB, FLT1*). **B:** Pathway analysis identifies GDF15 to increase expression of genes involved in inflammatory signaling (*TNFA, IFNγ, IL6*), angiogenesis (hypoxia, angiogenesis) and metabolism. **C:** Human primary HSCs (HHStec) were stimulated with either GDF15 (1μg/ml) or SPP1 (osteopontin, (1μg/ml)) for 24 hr and Western Blotting used to determine expression of IL-8 and phospho-STAT3. **D:** HSC (HHStec cells) were grown in monoculture, MP41 cells were grown in monoculture, HSC+MP41 cells were grown in co-culture, HSCs were treated with conditioned media (CM) from MP41 cells or GDF15 (1μg/ml) for 24 hr before probing for the expression of COL1A1 and PDGFRB by Western Blot. **E:** Immunofluorescence staining of COL1A1 in HSCs (HHStec) cells treated with basal media or conditioned media from MP41 or UM066 cells. **F:** CellChat analysis of human UM samples identifies HSC-derived collagens as being a potential outgoing signal to multiple immune subtypes and UM cells.

### UM-HSC derived factors increase vascular network formation

As many of the factors upregulated following co-culture of HSCs and UM cells were implicated in angiogenesis, we determined the role of GDF15 and IL-8 in vascular network formation. An analysis of likely interacting cell types from the CellChat analysis of primary and metastatic UM samples identified endothelial cells as a major target of secreted GDF15 (**Figure 6A**). CD4+, CD8+ T cells and NK cells were also predicted targets of GDF15. A human umbilical vein endothelial cell (HUVEC)-based vascular network assay demonstrated that both GDF15 and IL-8 increased the number and density of endothelial cell networks (**Figure 6B,C**). As both GDF15 and IL-8 are secreted by UM cells with levels increased in UM co-cultured with HSC we next treated endothelial cell cultures with conditioned media from UM/HSC co-cultures. As predicted, vascular network formation increased after treatment with conditioned media from MP41, UMM061or MM28 cells co-cultured with primary HSCs (**Figure 6D,E**). As the conditioned media was likely to contain other potential angiogenic growth factors we repeated the experiment in the absence and presence of a GDF15 blocking antibody (**Figure 6F,G**).The antibody prevented conditioned media-driven vascular network formation, indicating a key role for secreted GDF15 in the angiogenic response (**Figure 6F,G)**.

**Figure 6:**
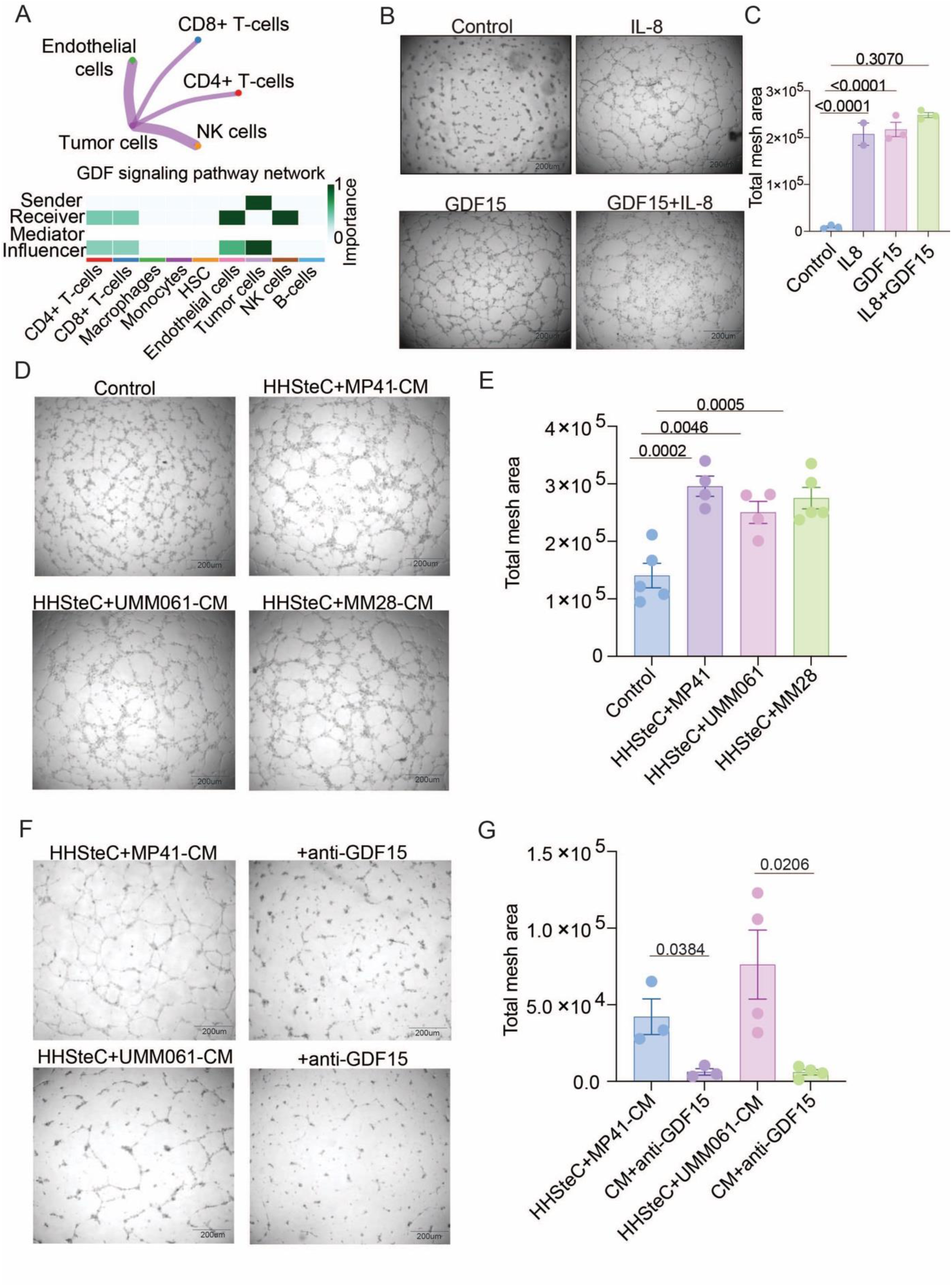
Both GDF15 and IL-8 drive vascular network formation. **A:** CellChat analysis of scRNA-Seq from human UM identifies endothelial cells as being a major target of secreted GDF15. **B:** Exogenous GDF15 and IL8 increase vascular network formation. HUVEC cells were plated on Matrigel and treated with basal media, IL-8 or GDF15 (both 1μg/ml for 0-24 hr). **C:** Scoring of endothelial network formation data from B. **D:** Conditioned media (CM) from UM-HSC co-cultures increases vascular network formation. HUVECs were plated onto Matrigel and treated with basal media or CM from MP41, UM066 and MM28 cells grown in co-culture with LX2 cells. **E:** Vascular network formation from D was quantified. **F:** GDF15-blocking antibodies reverse the pro-angiogenic effects of CM. HUVEC cells were treated with CM from MP41-LX2 co-cultures in the absence and presence of the GDF15 blocking antibody (Ponsegromab,120ng/ml) and vascular network formation imaged. **G**: Quantification of endothelial cell network formation from F.

### UM-derived GDF15 is required to establish liver metastases *in vivo*

scRNA-Seq data indicated that metastatic UM cells expressed high levels of GDF15 and that this increased HSC activation and angiogenesis. To evaluate the role of GDF15 in UM metastasis, we generated a mouse model of UM liver metastasis in which MP41 cells harboring a control (non-targeting) shRNA or a vector expressing an shRNA directed against GDF15 knockdown (**Supplemental Figure 2**) were injected into the tail vein (**Figure 7A,B: Supplemental Figure 3**). All of the mice with injected with cells containing the control shRNA developed liver metastases, whereas those injected with UM cells silenced for GDF15 did not develop metastases (**Figure 7A,B**). As it was difficult to determine whether UM cells depleted for GDF15 even reached the liver we developed a second model in which UM cells were injected by intraocular route (**Supplemental Figure 4**). After 1 week the injected eye was enucleated, and the development of metastases was tracked using IVIS imaging, with mean time to liver metastasis development being noted as 7 weeks. At endpoint, large macrometastases were observed on the surface of livers in the shRNA control group with no visible metastases seen in the GDF15 shRNA group. Histological examination of livers from both groups revealed the UM cells did reach the liver. In animals injected with UM cells with the GDF15 shRNA there were fewer, and much smaller, liver metastases than in mice injected with control shRNA UM cells (**Figure 7C,D**). The few liver specimens harboring GDF15-silenced UM metastases displayed reduced expression of HSC-associated ECM proteins such as fibronectin and Collagen 1A1(**Figure 7E-H**). The GDF15-silenced samples also showed a reduction in Ki67 staining relative to control tumors, suggesting that GDF15 was required for the fitness of UM liver metastases *in vivo* (**Figure 7I-J**). By contrast, silencing of GDF15 did not affect the rate of growth of MP41 cells *in vitro* (**Supplemental Figure 5**). Further analyses identified a dramatic reduction in angiogenesis in the GDF15 silenced tumors, as demonstrated by reduced CD31 staining (**Figure 7K,L**). Together, these findings point to a key role for UM-derived GDF15 in modulating the liver microenvironment, allowing for metastatic outgrowth.

**Figure 7:**
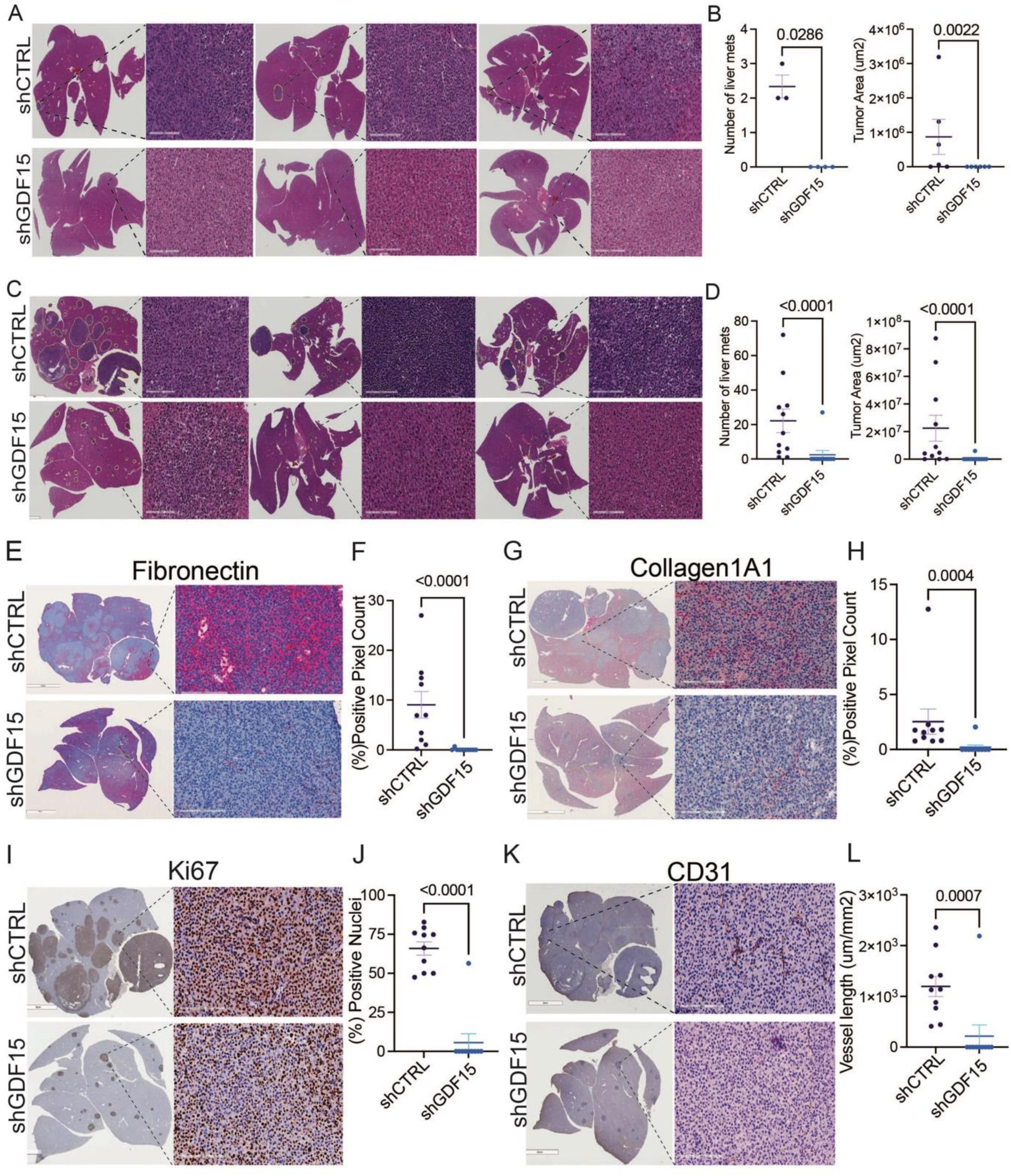
Silencing of GDF15 reduces the formation of UM liver metastases. **A:** H&E staining of livers following tail vein injection of GDF15 expressing and GDF15 silenced MP41 UM cells. Livers were collected after 6 weeks. **B:** Quantification of liver metastases following tail vein injection of GDF15-expressing or silenced MP41 cells. **C:** H&E staining of livers from the eye to liver metastasis model, demonstrating fewer and smaller liver lesions following GDF15 shRNA silencing. **D:** Quantification of the size and number of liver metastases from C. **E:** IHC staining of liver metastases for fibronectin in MP41 tumors either expressing or with GDF15 silenced. **F:** Quantification of data from E. **G:** IHC staining of liver metastases from MP41 tumors for the ECM protein Collagen 1A1. **H:** Quantification of data from G. **I:** IHC staining of liver metastases for Ki67 in MP41 tumors either expressing or with GDF15 silenced. **J:** Quantification of data from I. **K:** IHC staining of liver metastases for the endothelial cell marker CD31 in MP41 tumors either expressing or with GDF15 silenced. **L:** Quantification of data from K.

## Discussion

The remarkable propensity for uveal melanoma to metastasize to the liver ^1^ and the lack of effective treatment options for such patients^1^ has created a great need to understand the metastatic process and develop new interventions. Using high-dimensional single cell profiling to identify crosstalk among UM cells, HSCs and endothelial cells, we identified GDF15 as a key mediator of UM liver metastases.

scRNA-Seq analysis of UM monocultures and UM-HSC co-cultures demonstrated that HSCs modulated gene expression of the cancer cells, upregulating genes involved in survival such as *RasGRP3, ARF6* and mediators of AP-1 signaling, such as *FOS* and *JUNB* ^19, 20, 22^. RASGRP3 is a UM specific activator of Ras-driven MAPK signaling, which is activated through mutant GNAQ and PKC δ- and ε-dependent phosphorylation and diacyl glycerol (DAG) driven membrane recruitment ^19^. ARF6 is another small GTPase that acts as a central organizing node of GNAQ signaling in UM ^22^ that exerts signaling activity by trafficking GNAQ from the plasma membrane to cytoplasmic vesicles. Inhibition of ARF6 function can reduce UM tumorigenesis in cell culture and mouse models ^22^. In addition, co-culture of UM cells with HSCs also upregulated inhibitor of apoptosis (IAP) family member encoded by *BIRC7*, and *c-MYC*, also pro-survival factors ^20^. It thus appears that the interaction of UM cells with HSCs helps to tune oncogenic signaling from mutant GNAQ in the liver environment, increasing the likelihood of UM progression and survival. An analysis of scRNA-Seq data from clinical UM samples identified an increase in the strength of outgoing cell-cell interactions from the HSCs and endothelial cells in metastatic samples compared to the primary UM samples, suggesting these two host cell types are major microenvironmental drivers of liver metastatic disease. Further analysis identified a shared signaling signature between the UM cells and the HSCs in liver metastases which included growth factors previously implicated in UM liver metastasis such as HFG and FGF2 ^15, 16^. Another factor identified was PROS1, which stimulates the TAM (TYRO3, AXL, and MERTK) receptor tyrosine kinases. We previously reported increased PROS1 levels in metastatic UM, where it plays a role in regulating M1 to M2 macrophage polarization in the tumor microenvironment ^23^.

One of the key growth factors secreted by metastatic UM cells was GDF15, a TGF-β family member implicated in both cancer cachexia and vascular development ^24, 25^. Under physiological conditions, most tissues express low levels of GDF15 which can increase dramatically in response to stress, inflammation and injury ^25, 26^. GDF15 levels also rise in response to bacterial and viral infection, where they contribute to metabolic adaptations to infection through increased hepatic triglyceride export ^26^. Cancer is frequently associated with high levels of circulating GDF15 which contributes to cancer cachexia ^25^. In cutaneous melanoma, high serum levels of GDF15 are associated with worse overall survival ^27^. UM patients with clinically detectable metastases also have higher serum levels of GDF15 than individuals without ^28^.

Analysis of clinical UM specimens and metastatic and non-metastatic UM cell lines showed that elevated GDF15 expression is correlated with metastatic behavior. Metastatic dissemination in UM is frequently associated with loss of/mutation of BAP1 ^9^. Accordingly, shRNA-mediated depletion of BAP1 led to a strong increase in GDF15 expression, associated with increased H3K27ac at the GDF15 promoter. The increase in UM-derived GDF15 activated HSCs and endothelial cells, the two cell types predicted to interact strongly with UM cells in samples of human liver metastases. GDF15 was already known to play roles in liver inflammation and damage as carbon tetrachloride (CCL4) and chronic alcohol exposure increases hepatocyte GDF15 expression ^29^. While GDF15 was highly expressed in either BAP1 negative and/or PRAME-positive UM cells, low levels of GDF15 were also secreted from UM lines without these metastatic markers. scRNA-Seq and ELISA assays demonstrated that co-culture of nearly all the UM cell lines with HSCs increased GDF15 secretion, suggesting that HSCs amplify levels of GDF15 secreted by UM cells in the liver microenvironment.

GDF15 had multiple effects upon the differentiation state of HSCs including the upregulated expression of multiple ECM proteins, chemokines and growth factors, many including VEGF-A, IL-8, angiogenin and VCAM1 implicated in angiogenesis. The major GDF15-induced growth factor in HSCs was IL-8, an inflammatory mediator implicated in endothelial cell recruitment, vascular permeability and inflammation ^30^. There is already evidence that UM liver metastases exhibit increased levels of IL-8 expression and that *in vitro*, IL-8 plays a role in the radiotherapy-induced angiogenic response ^31^. In cutaneous melanoma, IL-8 is a well-established driver of metastatic development, in part through its effects on angiogenesis and through the release of pro-invasive matrix metalloproteinases such as MMP-2 ^32, 33^. It is significant that Class 2 UM cells did not secrete IL-8 and instead required crosstalk with HSCs to provide this pro-metastatic cytokine.

Co-culture and scRNA-Seq analyses suggested that the development of UM liver metastases involves close communication among UM cells, HSCs and endothelial cells. UM liver metastases are highly vascularized, and our *in vitro* studies suggest a role for both GDF15 and IL-8 in vascular network formation. In accordance with our findings, previous studies demonstrated that GDF15 increases endothelial cell proliferation through activation of the AP-1, MAPK and PI3K/AKT signaling pathways ^34^.

Silencing of GDF15 in UM cells inhibited the formation of large liver metastases but did not prevent the seeding of the cancer cells to the liver. Instead, metastases from GDF15-silenced UM cells demonstrated an impaired angiogenic response with very few infiltrating CD31+ endothelial cells and reduced levels of HSC elicited ECM proteins like Fibronectin and Collagen 1A1 that support tumor cell growth. Accordingly, while knock down of GDF15 did not affect UM cell growth, it significantly affected *in vivo* metastatic tumor proliferation, likely due to the impaired response of endothelial and HSC cells, resulting in very small metastatic lesions.

Although our focus was the effects of GDF15 on HSC differentiation and angiogenesis, it is likely that UM cell-derived GDF15 expression also impacts tumor immune recognition in the liver. GDF15 expression modulates LFA-1 expression on endothelial cells that represses the β2-integrin-mediated T-cell adhesion required for intravasation ^35^. High serum GDF15 levels in cutaneous melanoma and non-small cell lung cancer patients are predictive of failure to anti-PD-1 therapy ^35, 36^. Other potential immune effects of GDF15 include inhibition of T cell stimulation by dendritic cells ^37^ and the induction of regulatory T cell (Treg) activity during in hepatocellular carcinoma (HCC) leading to immunosuppression ^38^. Additional work has suggested that IL-8 can also drive immunosuppression in HCC through Treg polarization following the release of latent TGF-β ^39^. At this time, there are no therapies that can prevent the development of liver metastases in high-risk UM patients. Our results suggest use of a GDF15 blocking antibody, either alone or in combination with anti-PD-1, as in a clinical trial NCT04725474 for solid tumors could represent a novel therapeutic strategy in UM.

## MATERIALS AND METHODS

### Cell culture and reagents

RPMI culture medium was purchased from Corning (Corning, NY). Fetal bovine serum (FBS) was purchased from Sigma Chemical Co. (St. Louis, MO). The uveal melanoma cell lines 92.1, MP41, UMM055, UMM061, MM28, Mel202, Mel270, and Mel290 were obtained from Dr. J William Harbour (UT Southwestern Medical Center), OMM1 was provided by Dr. David Morse (Moffitt Cancer Center) and was used as previously described ^40^. Mel202 and 92.1 cells with inducible BAP1 silencing were described in ^23^. Human umbilical vein endothelial cells (HUVECs, #PCS-100-013) were acquired from ATCC (Manassas, Virginia). Human hepatic stellate cells (HSCs): LX2 (Cat. # SCC064) were obtained from Millipore Sigma and maintained with DMEM (#SLM-021-B) with 2% FBS and Pen/Strep. Primary human HSCs (HHSteC: #5300) were purchased from ScienceCell Research Laboratories and grown on poly-L-lysine-coated culture vessels (2 μg/cm2) in Stellate Cell Medium (#5301) with Stellate Cell Growth Supplement, 2% serum and Pen/Strep. All UM cell lines were cultured in RPMI-1640 supplemented with 10% FBS, L-glutamine, and antibiotics at 5% CO2. HUVEC cells were maintained in Endothelial Cell Growth Medium (Cell Applications Inc, San Diego, CA). Every 3 months, cells were tested for Mycoplasma contamination using the Plasmotest-Mycoplasma Detection Test (InvivoGen). Cell lines were authenticated by ATCC’s Human STR human cell line authentication service and cell lines were replaced from frozen stocks after 10 passages. Co-culture experiments were carried out under low (2%) serum concentrations.

### RNA Interference

shRNA viral particles targeting GDF15 (SHCCNV_NM-004864, TRCN000005839) were obtained from Millipore Sigma (MISSION® shRNA Lentiviral Transduction Particles. Multiple sequences were evaluated and the following sequence selected: (CCGGCCGGATACTCAGACCAGAAGTTCTCGAGAACTTCTGGTCTGAGTATCCGTTTTTG).

The non-targeting control (shControl) used was the MISSION® pLKO.1-puro Non-Target shRNA Control Transduction Particles (sequence: CCGGGCGCGATAGCGCTAATAATTTCTCAGAAATTATTAGCGCTATCGCGCTTTTT).

Infections were performed according to the manufacturer’s protocol. After 24 hours, the medium was replaced with fresh medium containing 0.5 µg/mL puromycin (Millipore Sigma) to select for successfully transduced cells. Polyclonal cells were plated as a single cell per well in a 96-well plate with a selection medium to isolate monoclonal cell populations.

### Single cell RNA-Seq on monoculture and co-cultures

A total of 5 × 10^5^ cells of LX2 HSCs were plated overnight, the following day 5 × 10^5^ UM cells (MP41) were added in HSC media and left to equilibrate overnight. After 24hrs, cells were replenished with fresh low serum media (2%). After 72hr of monoculture or co-culture, cells were processed for scRNA-Seq. Viable cells were loaded onto the 10X Genomics Chromium platform. The single cells, reagents, and barcodes were encapsulated into individual nanoliter-sized emulsions, and then the barcoded reverse transcription of poly-adenylated mRNA performed inside each individual encapsulation. Sequencing-ready cDNA libraries were completed in a single bulk reaction. RNA-Seq was performed on the Illumina NextSeq 500 using a high-output flow cell. Over 200,000 reads per cell were detected constituting around 1,031 genes per cell. De-multiplexing, barcode processing, alignment, and gene counting were performed on the 10X Genomics Cell Ranger 3.0 software.

### NicheNet

NichNetR was used on the scRNA-Seq data to infer cell-cell communications between the HSCs and UM cells ^41^. Gene expression in monoculture and co-culture conditions was compared to identify differentially expressed genes in the two conditions. We investigated which ligands from HSCs, as signal “senders”, potentially affected the expression of target genes in “receiver” tumor cells. Analysis followed a 5-step process: 1. HSCs were defined as the senders while tumor cells were defined as the receiver cells, and genes that were expressed in more than 10 percent in each cell type were filtered and kept for further analyses. 2. Differentially expressed genes in tumor cells between the co-culture and monoculture tumor samples were identified and defined as the gene set of interests in receiver cells. To narrow down the gene list, we further filtered retaining those that encode the receptors, highly expressed in > 30% of tumor cells when comparing co-culture-with monoculture conditions. 3. Ligands of interests were defined as those which are expressed in the HSCs and reported to bind a receptor expressed by the UM cells. 4. Potential ligands of interests were ranked based on the presence of their target genes in the gene set of interest (compared to the background set of genes). 5. Receptors and predicted target genes of interests in the ligand activity analysis were ranked.

### Cell-cell interaction analysis (CellChat)

CellChat analysis^42^ was performed on a previously published single-cell RNA sequencing analysis of primary and metastatic human UM samples ^21^ (GSE139829). To retain high-quality cells, we filtered and normalized the data to account for sequencing depth. To reduce the dimensionality of the dataset, we ran the function RunPCA() after scaling the data. Cell types were curated using BlueprintEncodeData. CellChat infers cell-cell communication based on the ligand-receptor signaling pathways involved between different types of cells. After identifying over expressed genes and interactions between the cells, we constructed aggregated the cell-cell communication network^42^. The communication network was visualized and plotted to analyze the number of interactions and the interaction strength of the signaling pathways between the different cell types.

### Processing of Published Data and Construction of the Core UVmap Reference for multiple scRNA-Seq datasets

In developing the core Uveal melanoma map (UVmap), we utilized the GBmap pipeline approach from Ruiz-Moreno et al. ^43^. We included only those samples confirmed as healthy eye or primary uveal melanoma, and healthy liver or metastatic liver, with each sample containing no fewer than 1,000 cells. A total of 6 datasets were included. The datasets for the core UVmap, which include Bakhoum et al.^44^, Lin et al.^45^, Ramachandran et al.^46^, and Gautam et al.^47^, were primarily in the form of raw count matrices. Where raw matrices were not available, BAM files were downloaded directly from the dbGaP cloud (Durante et al.^21^; phs001861.v1.p1; approved by dbGaP on May 24, 2022) or sourced directly from the authors (Pandiani et al.^48^), and were then transformed into FASTQ files and re-aligned using the STARsolo v2.7.10a pipeline (https://url.us.m.mimecastprotect.com/s/6yRCn5Gnyc0zDgOH9f3t1vGoW?domain=github.com). Gene names were updated to the most current HUGO nomenclature using HGNChelper ^49^, ensuring all clinical and diagnostic metadata remained consistent. Prior to integrating the datasets, stringent filtering parameters were applied to select only high-quality cells, excluding those with fewer than 500 genes, fewer than 1000 UMI counts (where applicable), and over 30% mitochondrial reads. Doublets in each droplet-based dataset were identified and removed using DoubletFinder ^50^. To mitigate batch effects across the datasets, a semi-supervised neural network model called single-cell ANnotation was employed using Variational Inference (scANVI) from Xu et al ^51^, within the transfer-learning framework of the single-cell architectural surgery algorithm (scArches) from Lotfollahi et al.^52^. scArches-SCANVI necessitates prior knowledge of cell types/labels for reference map creation. To standardize cell type labels from different sources, we annotated each dataset employing both automated and manual methods. For the automated process, we collected lists of melanoma and GEP markers from Durante et al. ^21^, 16 cancer cell states from Barkley et al.^53^, and a list of 174 adult eye and liver markers from a study published in Quan et al. ^54^. We then performed UCell signature scoring as per Andreatta et al.^55^ and applied a cutoff value of 0.2 to assign cells as state/marker-positive. Subsequently, manual cell identity was assigned based on results from the automated process, available original cell labels, and specific gene expression patterns analyzed via the Wilcoxon rank-sum test. CNV analysis was conducted using the CopyKAT package ^56^, categorizing cells as either diploid or aneuploid. This preliminary coarse cell type labeling facilitated the training and integration of the model through scANVI-scArches. The analysis was executed on the raw counts from the 5000 most variable genes, considering studies as the batch variable and adhering to recommended tool parameters. The output from the pipeline was a latent representation of the integrated data, which then served as input for clustering and dimensional reduction visualizations. Leiden clustering based on a k-nearest neighbor graph (k-NNG) ^57^ was applied to identify distinct cell populations, and Uniform Manifold Approximation and Projection (UMAP)^58^ for data embedding and two-dimensional reduction, using the plot1cell package ^59^ for UMAP visualization. Post-co-embedding, cell identities were refined manually for each cluster, utilizing our unified preliminary annotations and evaluating specific marker gene expression to accurately define each broad cell type/state.

### Cytokine arrays

The Human XL Cytokine Array from R&D Systems was used according to the manufacturer’s instructions (R&D systems, Minneapolis, Catalog# ARY022). Briefly, 5 × 10^5^ cells of HSCs (LX2) were plated overnight in low serum medium. Next day, 5 × 10^5^ UM cells (MP41or UMM061) were plated and allowed to equilibrate overnight. After 72 hr of monoculture or co-culture cells were trypsinized and whole cell lysates were extracted. Following BCA protein quantification, 200 μg of protein from mono or co-cultures was used for the cytokine arrays according to the manufacturers protocol. The images were quantified using ImageJ software. All the experiments were carried out 3 independent times.

### ELISA

Quantikine ELISA (R&D systems, Minneapolis, Catalog# D8000C, #DGD150) assays were carried out according to the manufacturer’s protocol. A total of 5 × 10^6^ HSCs (HHSteC) were plated overnight, followed, next day, by the addition of 5 × 10^5^ cells of UM cells (92.1, UMM055, Mel202, Mel290, UMM061, MM28, Mel270). After 72 hr of monoculture or co-culture supernatants were collected and analyzed as previously described ^60^. Optical densities were determined by measuring the wavelength at 540nm. Each condition was analyzed in duplicate in 3 independent experiments.

### Immunoblotting

Immunoblotting was carried out as previously described ^40^. The anti-phospho STAT3 (#9145), total STAT3 (#9139), IL8 (#94407), Collagen 1a (#39952), were purchased from Cell Signaling Technologies (Danvers, MA). BAP1 (#sc-28383, Santa Cruz Biotechnology, Dallas, TX, USA), GDF15 (# AF957, R&D systems). Anti-vinculin (V9131), anti-GAPDH (G9545) antibodies were from Millipore Sigma. All primary antibodies were used at a 1:1000 dilution. For recombinant proteins: human osteopontin (SPP1) (Sino biological #10352-H08H), GDF-15 (#957-GD, R&D Systems) were used at 1μg/ml to stimulate serum-starved HSCs. To induce the BAP1 silencing, both Mel202 and 92.1 cells (derived in ^23^) were treated with 1 μg/ml of doxycycline for 5 days before the collection of protein and immunoblotting as above.

### Chromatin Immunoprecipitation Sequencing (ChIP-seq)

ChIP-seq was carried out as previously described^61^. Briefly, 20 million cells were crosslinked for 7 min with 1% formaldehyde per experiment. The chromatin was sonicated to an average fragment size of 200–500 base pairs. For each ChIP experiment, 10 μg of antibody was used. The NEBNext Ultra 2 kit was used to prepare the libraries sequenced at a depth of >20 million reads per sample at the University of Miami Oncogenomics Shared Resource. The sequencing reads were filtered for quality and then aligned to the hg38 genome using Bowtie. Using MACS2, normalized coverage tracks were generated and plotted using SparK ^62^.

### Bulk RNA-Seq

3 × 10^5^ LX2 HSCcells were plated washed with PBS and incubated in serum-free media for 24hrs, before being treated with GDF15 at 1μg/ml for 24hrs. Total RNA was extracted using Qiagen Rneasy Plus Universal Mini kit following manufacturer’s instructions (Qiagen, Hilden, Germany). RNA sequencing libraries were prepared using the NEBNext Ultra RNA Library Prep Kit for Illumina using manufacturer’s instructions (NEB, Ipswich, MA, USA). Raw sequence data (.bcl files) generated from Illumina NovaSeq was converted into fastq files and de-multiplexed using Illumina bcl2fastq 2.20 software. Trimmed reads were mapped to the reference genome available on ENSEMBL using the STAR aligner v.2.5.2b. Unique gene hit counts that fell within exon regions were calculated by using feature Counts from the Subread package v.1.5.2.

### Endothelial tube formation assay

Tube formation assays were carried out as previous described^63^. The HUVEC cells (all below passage 5) were serum-starved for 24 hours before being plated on 50 μL of growth factor-reduced Matrigel. To check the effectiveness of GDF15, IL-8 or conditioned media (collected from HSC and UM cells) on tube formation, cells were plated with either basal media or 1μg/ml GDF15, IL-8, or UM-HSC conditioned media for 6 hrs. Images were captured to cover whole the field using the EVOS cell imaging system. The analysis of tube formation was carried as previously described ^64^. The effectiveness of blocking GDF15 on tube formation was assessed using an anti-GDF15 blocking antibody (ponsegromab: 120ng/ml, #HY-P99241, MedChemExpress, NJ, USA).

### Confocal microscopy

Immunofluorescence staining was carried out as previously described^65^. HSCs (7 × 10^4^) (HHSteC) were seeded on coverslips and serum-starved for 24hrs before being treated with conditioned media. 24hrs after treatment with conditioned media cells were fixed with 4% paraformaldehyde, permeabilized with 0.1% Triton X-100 for 10 minutes, and incubated with 5% bovine serum albumin (BSA). Cells were then labeled for 1 hour with primary antibody Collagen 1a (#39952, Cell Signaling Technology). After nuclear staining with DAPI (Sigma), the cells were visualized with a Leica SP5 confocal microscope.

### Liver metastatic models: Tail vein

All animal experiments were carried out in agreement with ethical regulations and protocols approved by the University of South Florida Institutional Animal Care and by The Institutional Animal Care and Use Committee (IACUC number IS000011179R). 8 weeks old female NSG mice from Jackson Laboratory were used (NOD.Cg-Prkdcscid Il2rgtm1Wjl/SzJ - Stock 005557). Each mouse was injected with 2×10^5^ MP41 shCTRL or shGDF15 MP41 UM cells into the tail vein. The tumors were allowed to grow for 6 weeks, before the mice were euthanized. The lungs and liver were collected, fixed and paraffin embedded. Sections were stained with hematoxylin and eosin (H&E). Slides were subsequently imaged and quantified the number of metastases using Aperio Imagescope.

### Suprachoroidal injections for liver metastases

1×10^5^ MP41 shControl or shGDF15 cells in 2μl of PBS were injected into the right eyes of sedated mice using a 33-guage Hamilton needle/micro syringe. 4 weeks after injection, the right eye was enucleated under anesthesia. The mice were observed for 17 weeks post enucleation before being euthanized. The livers, lungs, spleen, kidney hearth, colon and brain were collected and fixed in formalin for 48hrs followed by storage in 70% ethanol. The collected organs were paraffin embedded. stained with hematoxylin and eosin and subjected to immunohistochemistry (IHC) with an anti-CD31(Abcam, Catalog #ab28364), Ki67 (Abcam, ab16667) antibody. Slides were subsequently imaged and positive staining were quantified using Aperio Imagescope (Leica Biosystems, Richmond, IL).

### Statistical analysis

Results are expressed as mean ± standard error of the mean of at least three independent experiments. For the ELISA assays and tube formation experiments, one-way analysis of variance (ANOVA) was performed followed by a Sidak’s multiple comparison test for post-hoc analysis with non-significance equaling p > 0.05 and *=p < 0.05, **=p < 0.01, and ***=p < 0.001. Bulk RNA seq data analyses were carried out by log_2_Fold change and significant genes were calculated by Wald test and p adjusted by Benjamini-Hochberg *=p < 0.05. For the *in vivo* metastatic mouse models and anti-GDF15 studies significance were analyzed using non-parametric paired t test with significance equaling *=p < 0.05, **=p < 0.01, and ***=p < 0.001.

