## Supplementary figures and images for "GDF15 reprograms the microenvironment to drive the development of uveal melanoma liver metastases"

### Supplemental Figures 1-5

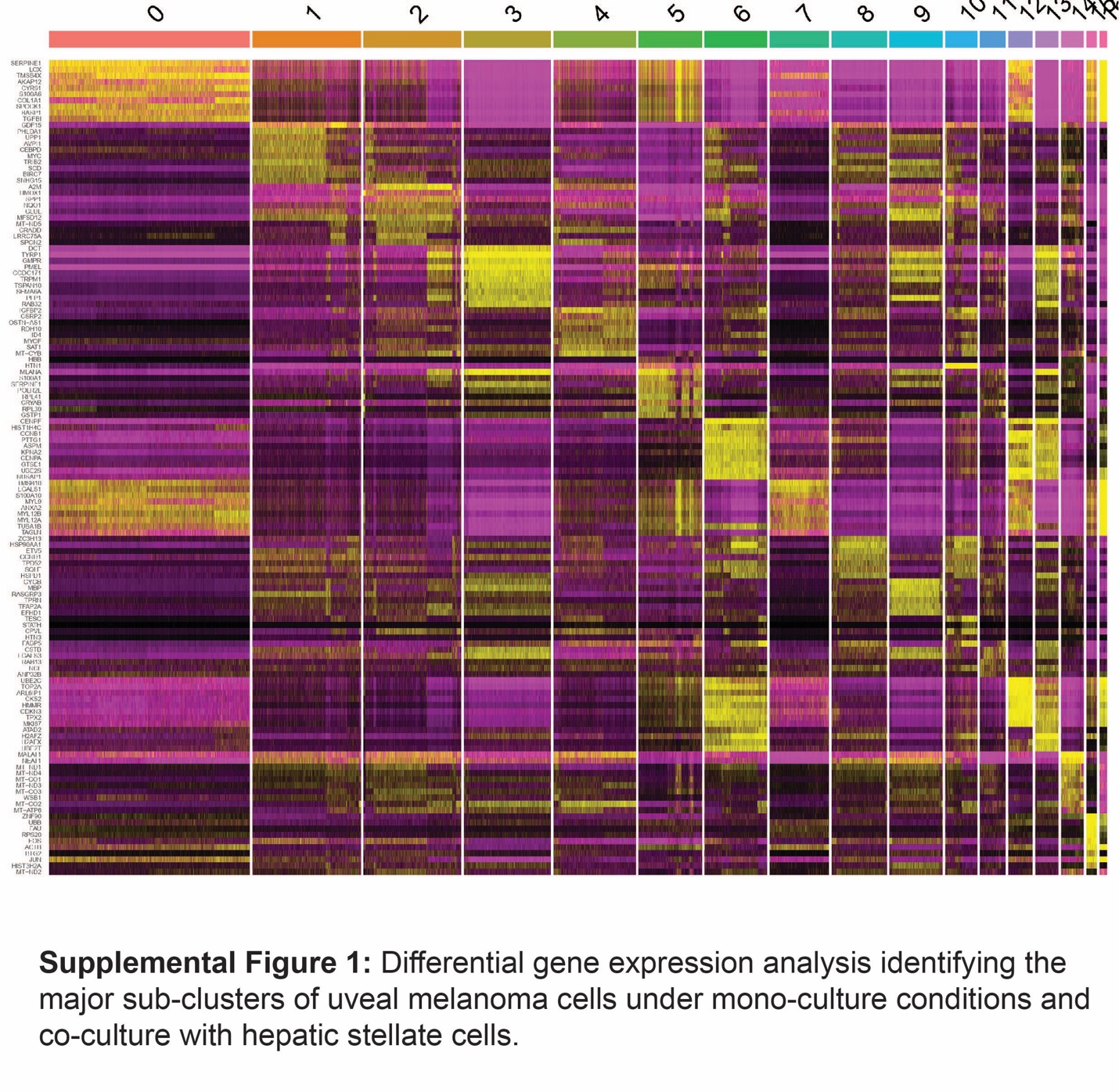

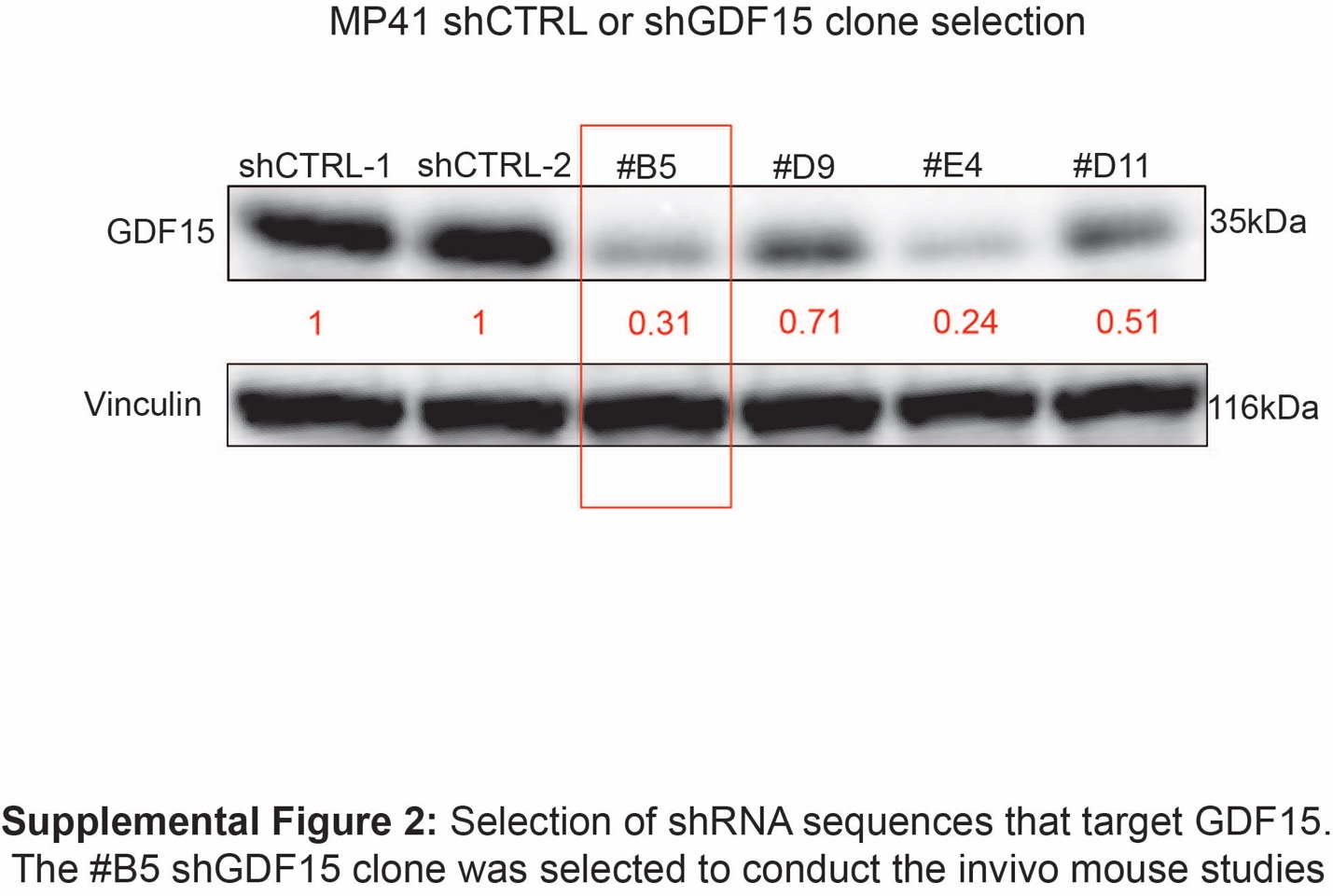

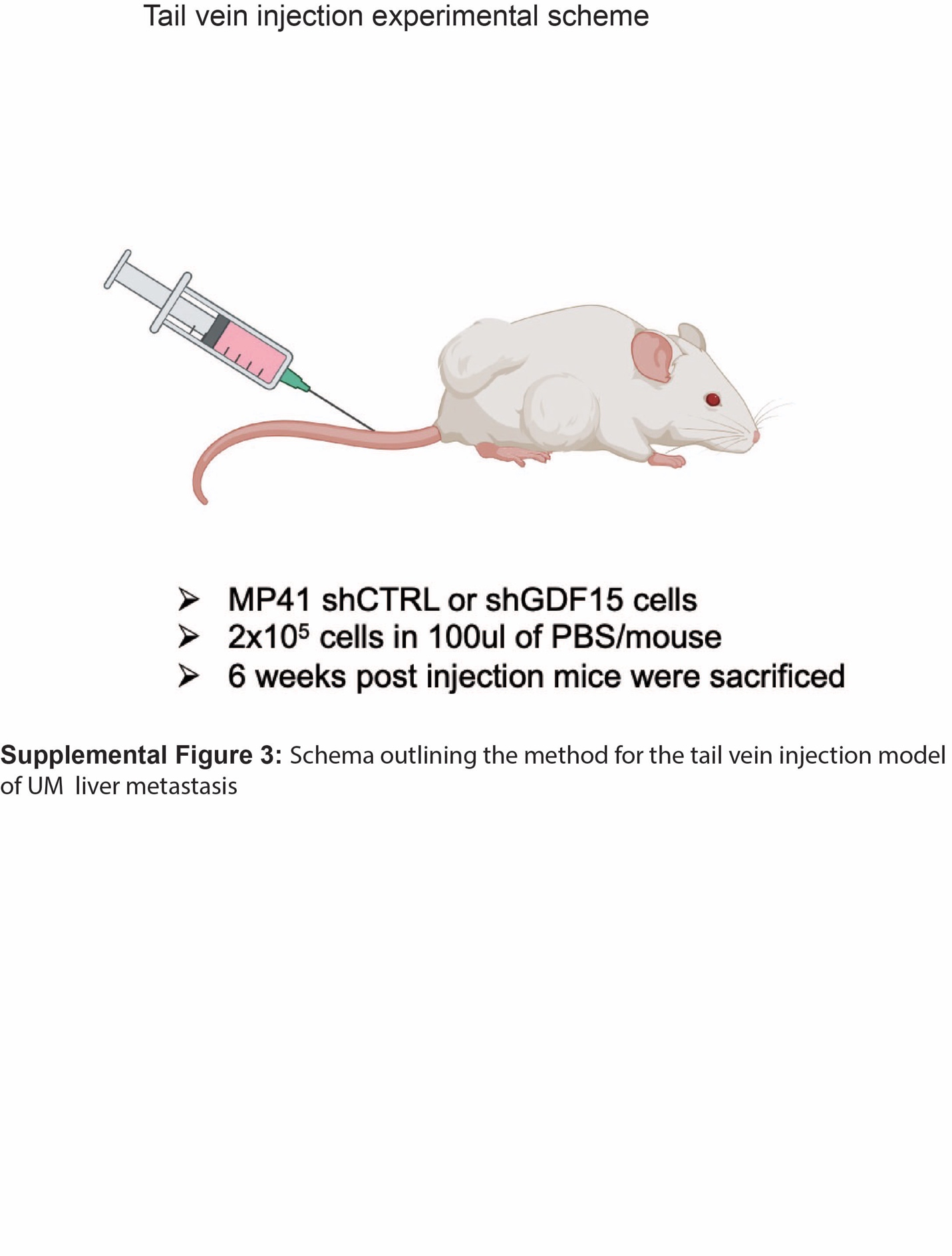

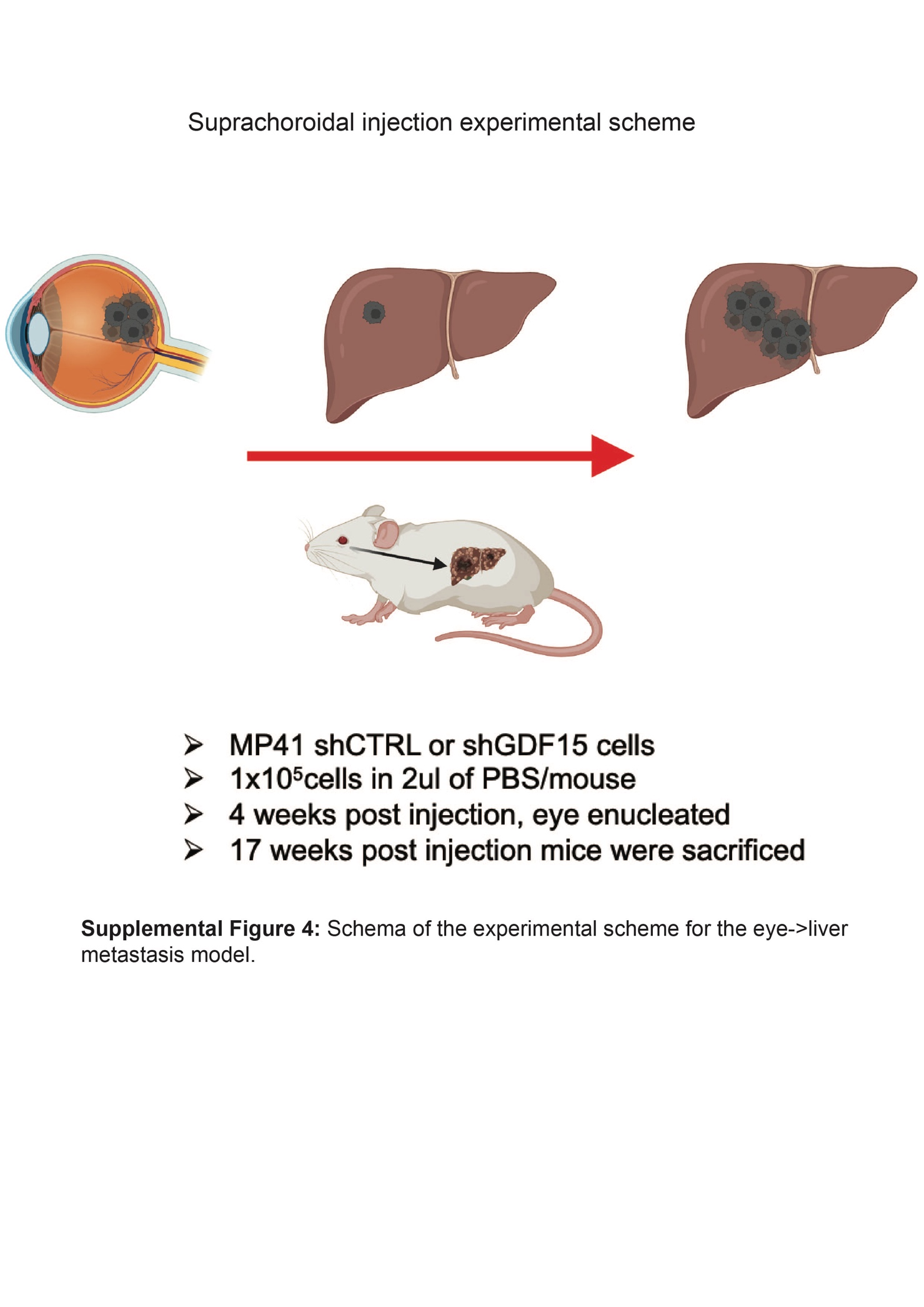

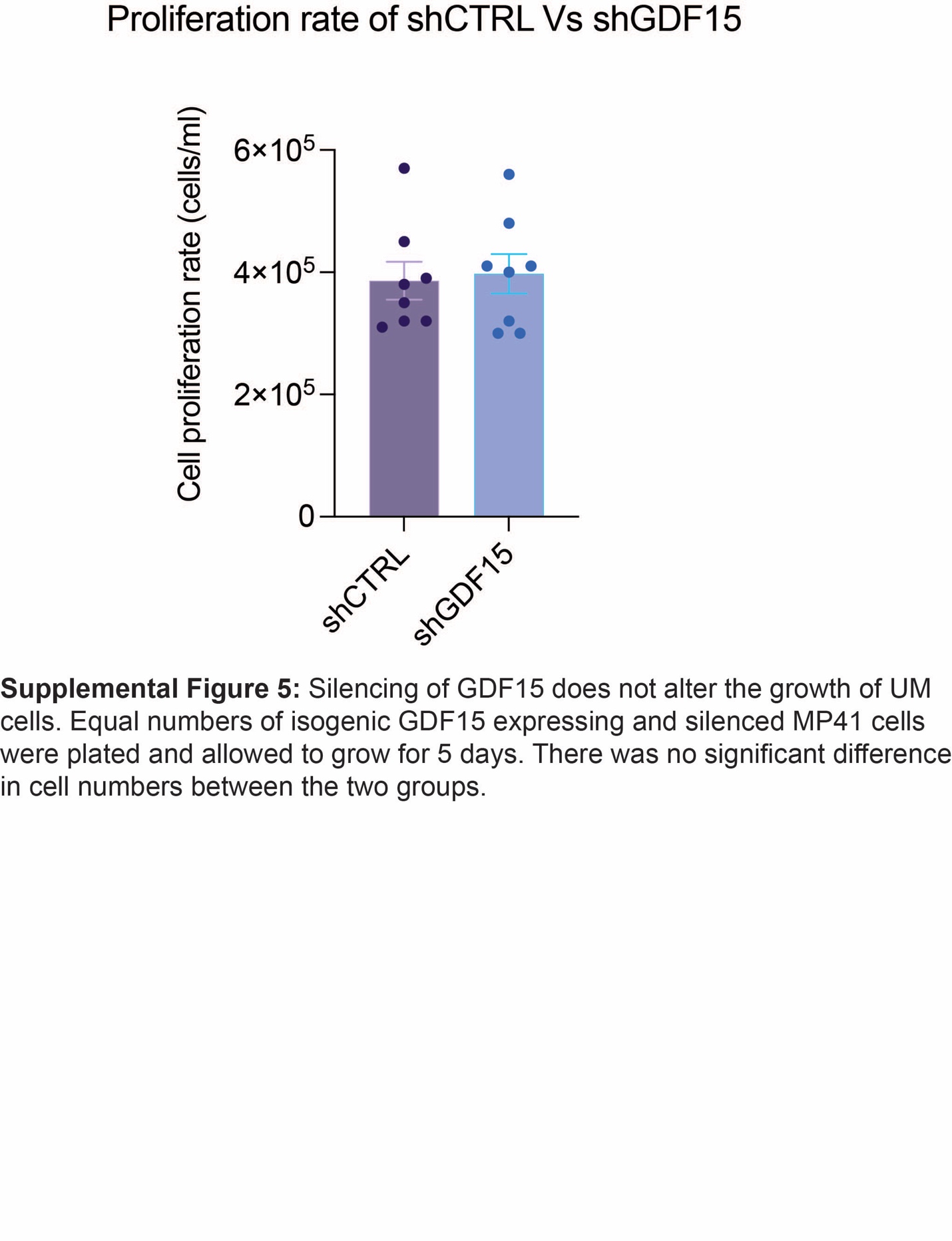
